# Inhibition of Replication Origins and ATR Synergistically Activates the Innate Immune System in Cancer Cells

**DOI:** 10.1101/2023.09.08.556350

**Authors:** Kai Doberstein, Johannes Panther, Sebastian Berlit, Benjamin Tuschy, Marc Sütterlin, Frederik Marmé

**Affiliations:** Department of Obstetrics and Gynaecology, Medical Faculty Mannheim of the Heidelberg University, University Medical Centre Mannheim, 68167 Mannheim, Germany; Mannheim Institute for Innate Immunoscience, Medical Faculty Mannheim of the Heidelberg University, 68167 Mannheim, Germany; DKFZ-Hector Cancer Institute at University Medical Center Mannheim, Mannheim, Germany

## Abstract

Sufficient numbers of activated replication origins are essential for successful DNA replication. While normal cells load replication origins in abundance to the genome, cancer cells often load fewer origins of replication due to genomic alterations such as *CCNE1* amplifications.

Here we exploit this feature to sensitize cancer cells to ATR inhibition.

We show by using siRNA and small molecules that the inhibition of origin activation as well as the reduction of replication origins itself sensitizes ovarian cancer cell lines against ATR inhibition by inducing genomic instability. We further show that in ovarian cancer cells this combinatorial approach leads to an increased genomic instability which manifests in an increase of micronuclei that consequently activate the innate immune system through the cGAS-STING pathway. Notably, no activation of the innate immune system was observed in immortalized fallopian tube cells. However, overexpression of Cyclin E1 in this model leads to a marked increase in genomic instability and innate immune activation.

Here we provide preclinical evidence to increase the therapeutic efficacy of ATR inhibitors. Additionally, this approach could help to activate innate immunity and increase T-cell immune infiltration in tumors, providing a rationale for a combination with immune checkpoint inhibitors.

## Introduction

Cancer cells rely on genomic instability to facilitate adaptation and evolution in response to changing environments and therapies. However, excessive levels of instability can be detrimental to cancer cells, underscoring the double-edged nature of this phenomenon. The inactivation of various DNA repair mechanisms can cause genomic instability, leading to diverse types of genetic aberrations, such as mutational burden, structural chromosomal alterations, and changes in ploidy.^1,2^ Multiple approaches have shown that genomic instability can be utilized to target cancer cells through synthetic lethality.^3,4^ Such strategies include PARP inhibition, targeting cells with defects in the homologous recombination (HR) pathway, and G2 checkpoint abrogation though checkpoint kinase 1 (CHK1) inhibition, which targets cells with defects in the G1 checkpoint.^5^ Additionally, multiple synthetic lethalities have been identified for the use of ATR inhibitors in tumors with oncogenic RAS, TP53 deficiency or ATM deficiency.^6–8^

Approximately 50% of high-grade serous ovarian cancers (HGSOC) have defects in the HR pathway, making them highly sensitive to PARP inhibitors as well as platinum salts.^9^ In contrast, tumors with an active HR pathway do not respond well to PARP inhibitors. Within the group of HR proficient tumors, the *CCNE1* amplified cancers account for about 20% and are associated with a primary treatment failure and reduced survival.^10,11^ High expression of Cyclin E1 leads to an increase in double-strand breaks (DSBs) and genomic instability.^12^ One reason for the increased genomic instability is the shortening of the G1 phase of the cell cycle due to elevated Cyclin E1 levels throughout the cell cycle. As a consequence cells don’t have enough time to acquire the necessary resources for the doubling of the genome during S-phase, such as nucleotides and replication origin loading.^12^ Since loading of replication origins in cells is restricted to the G1 phase, *CCNE1* amplified cells have a reduced number of replication origins.^13^ The reduction in origin licensing has been shown to increase genomic instability.^14^

Normal cells only use 10% of their replication origins, and keep the rest of dormant origins as a reserve for replication stress.^15–17^ Cancer cells with a reduced number of dormant origins could therefore are more sensitive to a further reduction in replication origins loading as well as replication activation compared to normal cells.

The ATR kinase is important for the response to replication stress and replication fork problems ^18–20^ by supporting fork restart and dormant replication origin control. Importantly, ATR inhibitors lead to a strong activation of dormant replication origins and a reduction in replication speed.^21^ The activation of dormant replication origins ensures the safe duplication of the genome during S phase. Cancer cells with a reduced number of replication origins might therefore not be able to replicate successfully during S-phase of the cell cycle.

Based on the critical role of ATR kinase in the response to replication stress, we investigated whether cancer cells with a reduced number of replication origins may be more sensitive to ATR inhibitors. In this study, we examined the effects of reducing replication origins or origin activation in combination with ATR inhibition, using both siRNA and small molecule inhibitors. Our findings suggest that this combined treatment approach may offer a promising therapeutic strategy for targeting cancer cells with genomic instability and dysregulated cell cycle control.

## Results

### Sensitivity to ATR inhibitors negatively correlates with the expression replication origin genes

To analyze a potential association between the number of replication origins and the sensitivity to ATR inhibition, we compared the sensitivity (Area Under the Curve, AUC) to the ATR inhibitor AZD6738 (referred to as ATRi within this manuscript) of multiple ovarian cancer cell lines in the Cancer Cell Line Encyclopedia (CCLE) to the mRNA expression of genes involved in the replication origin *(MCM2-7, CDC6, CDT1* and *ORC1*). In parallel we also investigated the expression of *ATR* and *CCNE1*. As expected, the sensitivity to the inhibitor was associated with *ATR* expression, whereas the replication origin genes were separated in a second major cluster (blue cluster, Fig. 1a). Interestingly, we found a negative correlation between the expression levels of most replication origin genes and the sensitivity towards AZD6738, suggesting that cells that are more sensitive to ATR inhibitors are dependent on higher expression levels of replication origin genes (Figure 1b). The most pronounced negative correlation was observed between the ATRi and the expression levels of *ORC1* and *MCM5* (Figure 1b, c). To validate the generalizability of this effect beyond AZD6738, we investigated the impact of four additional ATR inhibitors (VE822, VE821, QL-VIII-58, and AZ20). Consistent with AZD6738, we found that the AUC and half-maximal inhibitory concentration (IC50) values of all four ATR inhibitors were predominantly inversely correlated with the expression of origin genes (Figure 1d). These findings suggest that cells exhibiting higher sensitivity to ATR inhibitors may rely on a larger number of replication origins for their survival.

**Figure 1:**
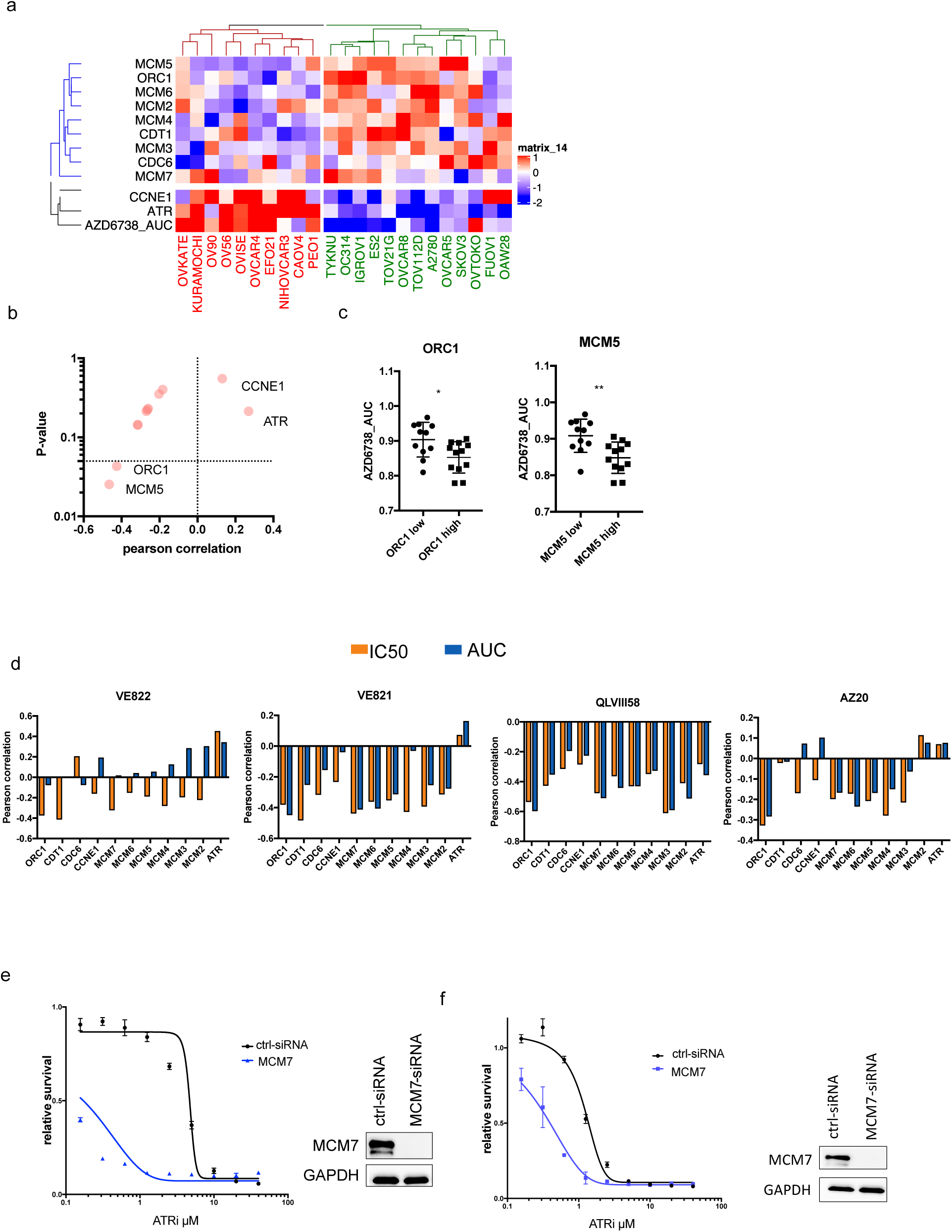
Reducing replication origins sensitizes cells to ATR inhibitors. a) Heatmap showing the RNA expression of different genes and the sensitivity to the ATR inhibitor AZD6738 in multiple ovarian cancer cell lines from the CCLE dataset. Cell lines were divided into an origin of replication high cluster (green) and an origin of replication low cluster (red). The gene expression of the origin of replication genes separated to the blue cluster in contrast to the black cluster that represents CCNE1 and ATR expression as well as the area under the curve (AUC, a low value represents more sensitive cells) of the ATR inhibitor AZD6738. b) vulcano blot showing the Pearson correlation and p-value between the RNA expression of genes and the sensitivity to AZD6738. c) Scatter plot of the expression of ORC1 (left) and MCM5 (right) against the sensitivity against AZD5738. d) Pearson correlation of genexpression against the IC50 or AUC of the ATR inhibitors VE822, VE821, QLVIII58 and AZ20. e and f) OVCAR8 (e) and OVSAHO (f) cells that were treated with siRNA against MCM7 or a control siRNA. 72 hours after siRNA transfection cells were treated with different concentrations of the ATR inhibitor AZD5738. Left depicts the relative survival and on the right is the knock down efficiency depicted and analyzed by western blot.

### Reducing replication origins sensitizes cells to ATR inhibitors

It has been shown that the inhibition of ATR activates origin firing.^21^ The increase in origin firing helps cancer cells during replication stress to secure genome integrity by completing genome duplication within S-phase. Based on this understanding, we hypothesized that further reducing the number of replication origins in cancer cells, might impair their ability to complete genome duplication and lead to intolerable levels of genomic instability during ATRi treatment. To test this, we analyzed if the downregulation of the minichromosome maintenance protein complex (MCM) might sensitize ovarian cancer cells to ATR inhibitors.

MCM7 along with MCM2-MCM6, forms during the G1 phase a heterohexamer that is required for both DNA replication initiation and elongation. Disruption of any MCM protein leads to the destabilization and degradation of the complex. We observed that the loss of MCM7 sensitized cells for the treatment with ATRi. In OVCAR8 cells, loss of MCM7 resulted in an almost 50-fold reduction in IC50, while in OVSAHO cells, a 4-fold reduction was observed (Figure 1e, f).

In summary, our findings demonstrate that reducing the expression of replication origin genes sensitizes ovarian cancer cells to ATRi treatment.

### Reducing replication origins by small molecule inhibitors

We next investigated whether compounds that reduce origin loading or origin activation could sensitize cells to ATR inhibitors. To accomplish this, we employed XL413 and RL5a, which have been shown to respectively inhibit replication origin activation and reduce the number of MCM proteins. RL5a achieves this by transcriptionally reducing the available MCM proteins, while XL413 inhibits CDC7 kinase, preventing the formation and activation of the Cdc45-MCM-GINS (CMG) complex and phosphorylation of MCM proteins.^22,23^ Recent studies have highlighted the cooperation between CDC7 and ATR in coordinating DNA synthesis with cell-cycle transitions using XL413 and ATRi.^24^ We first tested its ability to reduce origin of replications and origin activation (Figure 2 a, b). Consistent with previous findings, we observed with both treatments a reduction in phospho-MCM2 levels, while RL5a treatment also resulted in reduced levels of MCM2 and MCM7.

**Figure 2:**
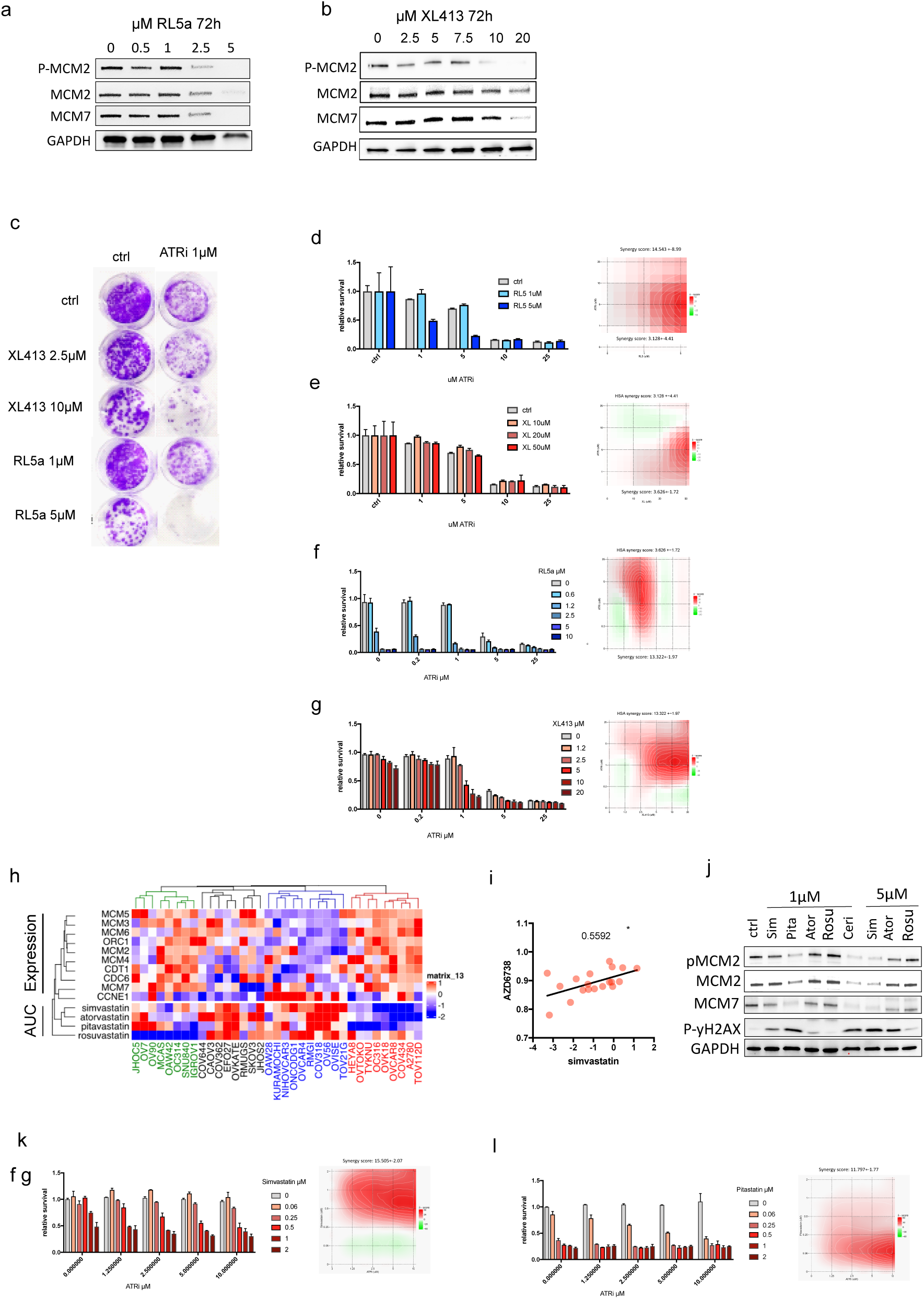
Compounds that reduce replication origins or origin activation work synergistic with ATR inhibitors. a and b) Western blot of OVCAR8 cells treated with different concentrations of RL5a (a) or XL413 (b). c) Colony formation assay of OVCAR8 cells, pretreated with XL413 or RL5a for 72 hours followed by the treatment with ATRi. d-g) Survival analysis of OVCAR8 (d, e) and OVSAHO cells (f, g) treated with different concentrations of RL5a (d, f) and XL413 (e, g) followed by ATRi treatment. On the right is the synergy profile depicted. h) Heatmap showing the RNA expression of different genes and the sensitivity, measured in area under the curve (AUC), to simvastatin, atorvastatin, pitavastatin and rosuvastatin in multiple ovarian cancer cell lines. i) Scatter plot with Pearson correlation (R=0.5592, *: p<0.05) of the sensitivity of AZD6738 and simvastatin in multiple ovarian cancer cell lines from the CCLE dataset. j) Western blot expression in OVCAR8 cells after the treatment with 1µM or 5µM of different statins. k and l) Survival analysis of OVCAR8 cells treated with different concentrations of simvastatin (k) or pitavastatin (l) followed by ATRi treatment. On the right is the synergy profile depicted.

Since MCM proteins are relatively stable and approximately 50% of MCMs are recycled from the previous cell generation, we pre-treated the cells with RL5a or XL413 for 3 days before adding the ATR inhibitor.^25^ In colony formation assay, we observed a synergistic effect in OVCAR8 cells when combining RL5a or XL413 with ATRi after pre-treatment (Figure 2c). Interestingly, when treating relatively dense cells, we only observed synergy with RL5a and not with XL413 in OVCAR8 cells (Figure 2d, e). Notably, in OVSAHO cells, we observed a stronger synergy with XL413 compared to RL5a (Figure 2f, g). In summary, our results demonstrate a synergistic effect between pre-treatment of XL413 or RL5a and ATR inhibitors in different cell lines, providing further evidence of their potential as a combinatorial approach.

### Statins can reduce the origin pool

Multiple studies have shown that statins are able to reduce the amount MCM mRNA and proteins in cells.^26,27^ We therefore investigated whether pretreating ovarian cancer cells with statins could sensitize the cells to ATR inhibitors. We compared the sensitivity of multiple cell lines to different statins, including Simvastatin, Atorvastatin, Pitavastatin, and Rosuvastatin, with the expression of genes involved in replication origin. Interestingly, the cell lines that were most sensitive to Simvastatin, Atorvastatin, and Pitavastatin exhibited the highest levels of replication origin genes (Figure 2h, red cluster). A separate panel of cell lines was sensitive against Rosuvastatin, showing also a high expression of origin genes (green cluster). Conversely, cell lines with the lowest expression of origin genes and the highest *CCNE1* expression were most resistant to various statins (blue cluster). Notably, Simvastatin showed the strongest negative correlation with the expression of origin genes (Sup. Figure 1a).

When comparing the sensitivity of ovarian cancer cell lines to ATRi and different statins, we found that, except for Atorvastatin, the sensitivity to all statins showed a positive correlation with the sensitivity to AZD6738, with the sensitivity to Simvastatin being the only statin significantly correlated with the the sensitivity to ATRi (Figure 2i, Sup. Figure 1b). These data indicate an association between the expression of genes involved in the replication origin process and the sensitivity towards statins as well as ATRi.

Ovarian cancer cell lines were most sensitive to Cerivastatin, followed by Pitavastatin and Simvastatin, while Atorvastatin and Rouvastatin showed the least effect on cell viability in different ovarian cancer cell lines (Sup. Figure 1c). Treatment with a concentration of 1µM reduced MCM2/7 protein expression with Simvastatin, Pitavastatin and Cerivastatin, but no effects were observed with Atorvastatin and Rosuvastatin (Figure 2j). Additionally, at a concentration of 5µM, we observed a reduction in MCM2/7 with Atorvastatin but not with Rosuvastatin. These findings are consistent with recently published microarray data in pancreatic carcinoma cells.^27^ Interestingly, the reduction of MCM2/7 was paralleled with an increase in phospho-yH2AX, indicating an increase in replication stress (Figure 2j). Based on the effectiveness to reduce MCM levels and the lower toxicity, in contrast to Cerivastatin, we decided to focus on Simvastatin and Pitavastatin for further studies (Sup. Figure 1c).

We found that pretreatment with 0.5 to 1 µM of Simvastatin sensitized OVCAR8 towards ATRi (synergy score: 15.5, Figure 2k). For Pitastatin a concentration of 0.06 µM was optimal to sensitize OVCAR8 to ATRi (synergy score: 11.8, Figure 2l). In OVSAHO cells, a concentration of 2 µM Simvastatin or 0.4 µM Pitavastatin was necessary to sensitize the cells for the ATR inhibitor AZD6738 leading to synergy scores of 6.7 or 7.4, respectively (Sup. Figure 1d, e, f). In summary, we demonstrate that repurposing drugs that reduce the number of replication origin loading can sensitize ovarian cancer cells to ATRi.

### Reduction of the replication origins leads to chromosomal abnormalities

We observed that the combined reduction of replication origins and treatment with ATR inhibitors resulted in a significant increase in the number of abnormally formed nuclei, indicating a rise in genomic instability (Figure 3a). While we did not observe synergistic changes in r-loop formation or p-yH2AX signal when combining the loss of replication origins with ATR inhibitors, we did observe alterations in the cell cycle, as reflected by changes in the CDT1/Geminin ratio (Sup. Figure 2a-c), which were further confirmed through cell cycle analysis (Sup. Figure 2d). By using fiber spread assay, we observed a considerable decrease in replication speed when treating MCM7-reduced cells with ATR inhibitors (Figure 3b). Studies have demonstrated that increased genomic instability can lead to the formation of micronuclei, which harbor sections of genomic DNA.^28^ The instability of micronuclei results in the exposure of double-stranded DNA to the cytoplasm. The cGAS-STING pathway, an evolutionarily conserved sensor of the innate immune system, can detect this DNA and activate innate immune signaling. The combined reduction of replication origins using *MCM7* siRNA and treatment with ATRi lead to a significant increase in cGAS expression and the number of cGAS-positive cells (Figure 3c, d). This effect was also accompanied by an increase in active P65 and STAT1, as well as an increase in the number of cells with micronuclei (Figure 3c, d).

**Figure 3:**
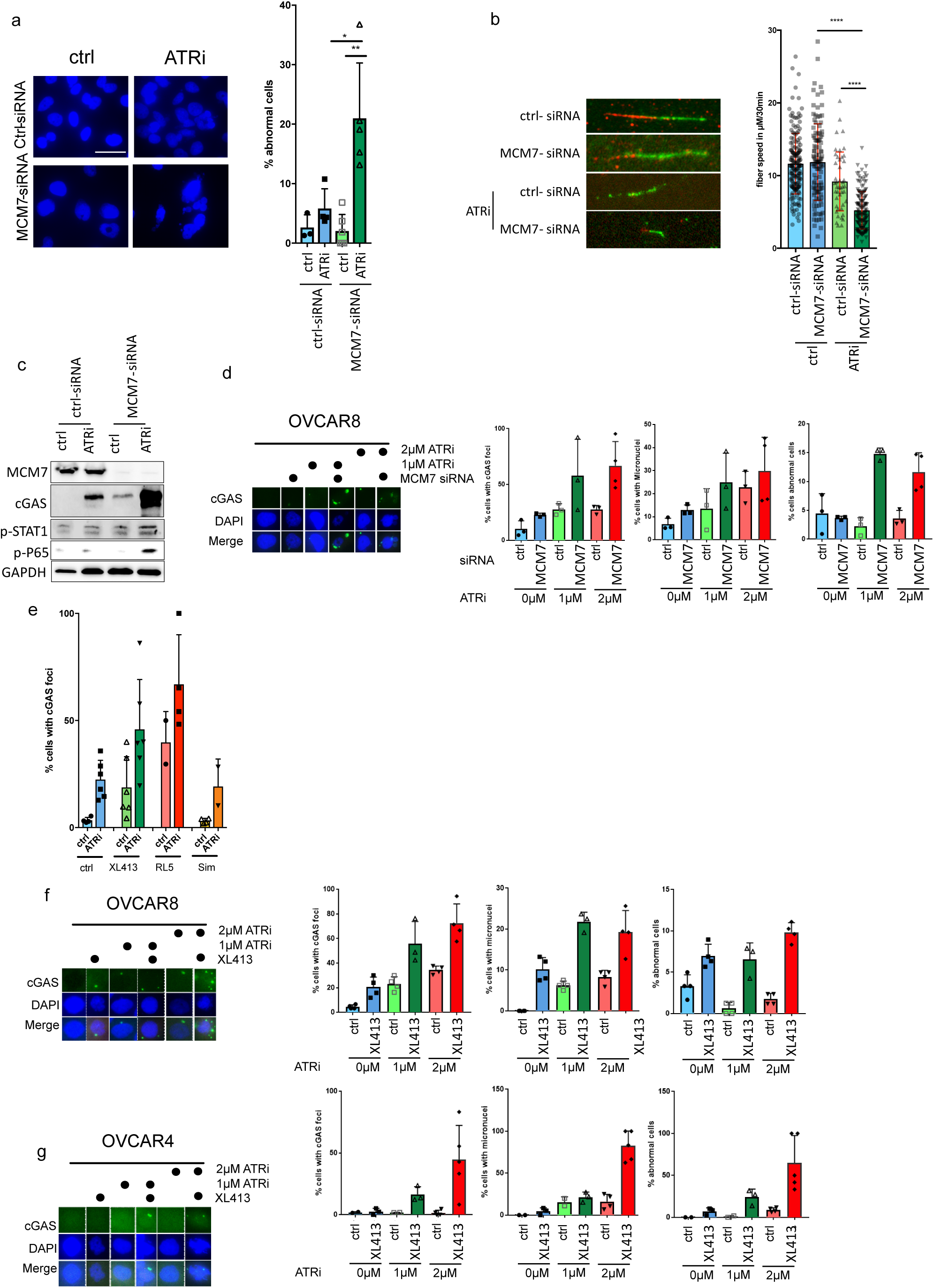
Reduction of the replication origins leads to chromosomal abnormalities. a) Left: Fluorescence DAPI analysis at 40x of nuclei that were treated with siRNA against MCM7 followed by ATRi treatment. Right: bar graph depicting percentage of abnormal cells analyzed in one picture. Scale bar 50 μm b) DNA fiber spread assay as described in material and methods. Left: representative picture of replication fibers under different conditions described in a (100x objective). Right: Fiber speed measured under different treatment conditions. c) Western blot analysis of OVCAR8 cells treated with siRNA against MCM7 followed by ATRi treatment. d) OVCAR 8 cells treated with control or MCM7 siRNA followed by the treatment with 1 or 2 µM ATRi. Left: representative immunofluorescence analysis showing nuclear DNA (DAPI, blue) and cGAS accumulation (green). Right: Immunofluorescence pictures were analyzed for % cells with cGAS foci, % cells with micronuclei and % abnormal cells. e) OVCAR8 cells pre-treated with XL413, RL5a or Simvastatin followed by the treatment of ATRi. Cells were analyzed by immunofluorescence for cGAS foci and percent cells with cGAS foci per image are depicted. f and g) OVCAR 8 (f) and OVCAR4 (g) cells treated with vehicle or XL413 followed by the treatment with 1 or 2 µM ATRi. Left: Immunofluorescence showing nuclear DNA (DAPI, blue) and cGAS accumulation (green). Right: Immunofluorescence pictures were analyzed for % cells with cGAS foci, % cells with micronuclei and % abnormal cells.

We further investigated whether the same effects could be observed when combining ATR inhibitors with XL413, RL5a, or Simvastatin. Compared to ATR inhibitor treatment alone, we observed an increase in cGAS expression when combining ATR inhibitors with XL413 or RL5a. However, no increases were observed when combining Simvastatin with ATR inhibitors compared to ATR inhibitor treatment alone (Figure 3e). Due to the lower activation observed with XL413 as a single treatment compared to RL5a, we decided to analyze XL413 more closely. The synergistic effects between XL413 and ATR inhibitors were observed in both OVCAR8 and OVCAR4 cells (Figure 3f, g).

In summary we could show that the reduction of replication origins or origin activation leads to genomic instability and activates the innate immune cGAS-STING pathway.

### cGAS expressing tumors are associated with a high replication origin gene expression

We aimed to identify tumor subgroups that might exhibit increased sensitivity to replication origin reduction and ATR inhibitors. For this purpose, we analyzed the TCGA ovarian cancer dataset. As expected, *cGAS* expression correlated with INF-gamma-response, TH2-cell infiltration and in multiple datasets with cytotoxic T-lymphocytes infiltration (Figure 4a, b, Sup. Figure 2e).

**Figure 4:**
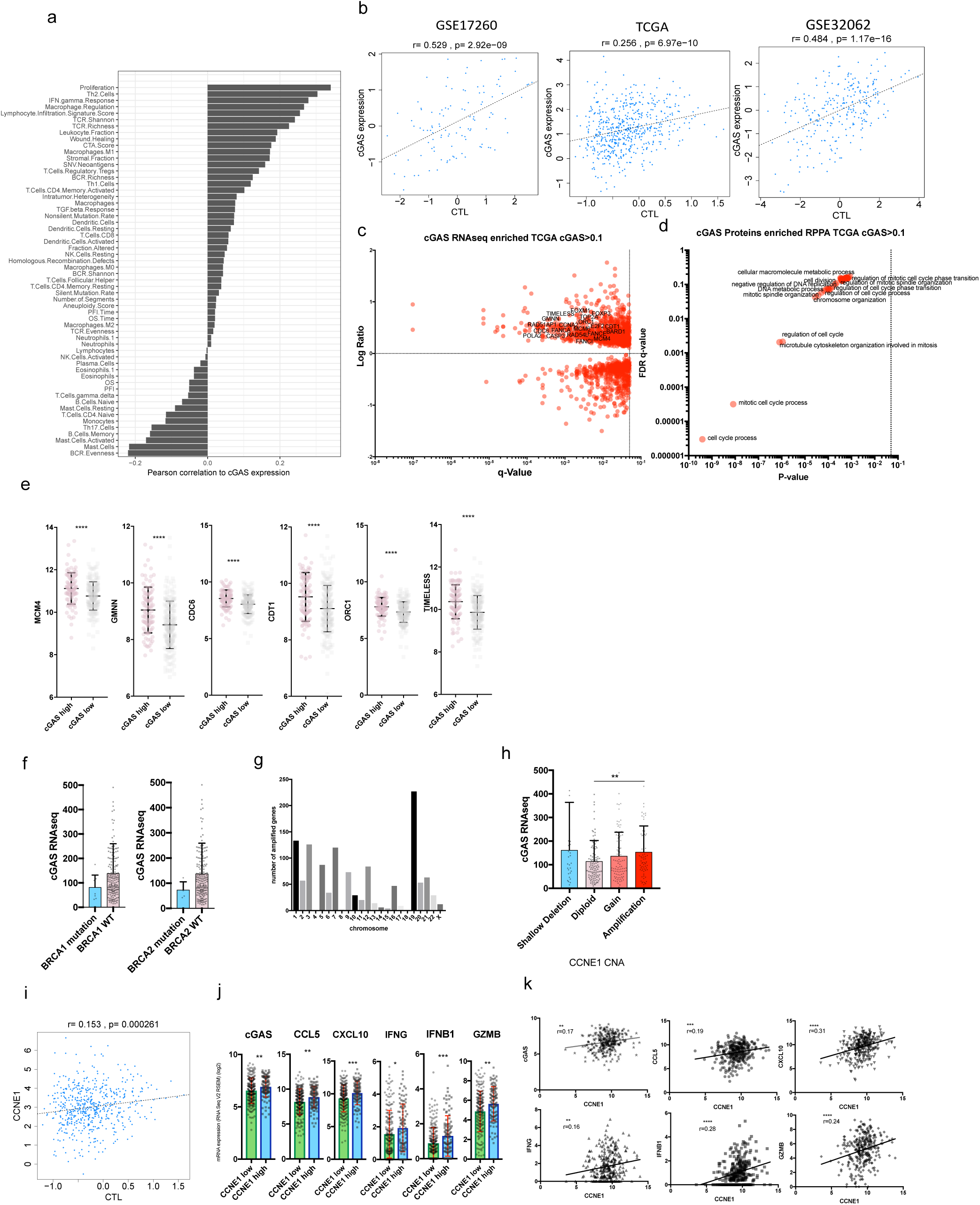
cGAS expressing tumors are associated with a high replication origin gene expression. a) Pearson correlation of the cGAS RNA expression of the TCGA ovarian cancer dataset against a panel of immune scores from Thorsson et al.^53^ b) Correlation of cGAS RNA expression in 3 different patient datasets against Cytotoxic T-cell (CTL) score from the TIDE project.^50^ c) Genes enriched in cGAS high expressing tumors from the TCGA ovarian cancer dataset (RNAseq cGAS expression >0.1). d) Enriched pathways from significantly enriched genes from c. e) Expression differences in cGAS high and cGAS low tumors of the TCGA ovarian cancer dataset. f) cGAS expression in BRCA1 (left) and BRCA2 (right) mutated tumors of the TCGA ovarian cancer dataset. g) number of amplified genes per chromosome in cGAS high expressing tumors analyzed in the TCGA dataset. h) cGAS RNAseq expression in ovarian cancer cells with different CCNE1 copy number status analyzed in the TCGA dataset. i) Correlation of CCNE1 RNA expression in patients against the CTL score in the TCGA ovarian cancer dataset. j) RNA expression of different genes in CCNE1 high and CCNE1 low tumors analyzed in the TCGA dataset. k) RNA Correlation of different genes with CCNE1 expression in the TCGA ovarian cancer dataset.

Pathway enrichment analysis revealed that cell cycle progression and mitotic cell cycle progression were the most significantly enriched pathways (Figure 4c, d, Sup. Figure 2f,g). This finding aligns with recent research from the Greenberg lab, which demonstrated that cell cycle progression is necessary for the occurrence of micronuclei and cytoplasmic DNA during radiation therapy, subsequently leading to activation of the cGAS-STING pathway.^29^

The enriched genes included replication origin licensing genes such as *MCMs*, *CDC6*, *CDT1*, *ORC1*, as well as cell cycle regulation and mitosis-related genes such as *GMNN* and *TIMELESS*, indicating a connection between these pathways (Figure 4c, e). Since HGSOC is a copy number driven cancer, there are only a few recurrent mutations. Two of those are the *BRCA1* and *BRCA2* genes that play an essential role during HR DSB repair. While *cGAS* expression showed a reduction in *BRCA1* and *BRCA2* mutated tumors, it did not reach statistical significance (Figure 4f). We also did not observe an association between HRD score and *cGAS* expression (Sup. Figure 2h). When analyzing the location of amplified genes that were enriched in *cGAS* high expressing tumors, we found that the chromosome harboring most genes was chromosome 19 (Figure 4g). Notably, chromosome 19 is also the location of *CCNE1*, which is frequently amplified in various tumor types, including in approximately 20% of HGSOCs. Cyclin E1 overexpression results in the shortening of the G1 phase, limiting the time available for replication origin loading during this phase and leading to a reduced number of replication origins. Furthermore, previous studies have shown that high Cyclin E1 levels can interfere with replication loading.^13,30^ *CCNE1* amplified tumors express significantly more cGAS than diploid tumors. Interestingly, tumors with a shallow *CCNE1* deletion also express more *cGAS* compared to diploid cells (Figure 4h). Consistent with this, *CCNE1* expression weakly but significantly correlated with cytotoxic T-cell infiltration, as well as increased expression of *cGAS*, *CCL5*, *CXCL10*, *IFNG*, *IFNB1*, and *GZMB* (Figure 4i-k). Therefore, we speculate that Cyclin E1 overexpression might contribute to cGAS activation.

In summary, our findings demonstrate that cGAS plays an important role in immune tumor infiltration and its expression is strongly connected to the expression of cell cycle and replication genes.

### Reducing the replication origin pool in *BRCA2* mutated cells

We aimed to investigate whether *BRCA* mutations affect the synergistic effects between the reduction of replication origins and ATR inhibition in ovarian cancer cells. Approximately 50 percent of ovarian cancer tumors exhibit defective homologous recombination (HR) DNA repair pathway, commonly caused by somatic or germline mutations in HR genes such as *BRCA1* or *BRCA2*. To explore this, we generated OVCAR8 clones with a defective *BRCA2* gene using CRISPR/CAS9 (Sup. Figure 3a). These clones showed loss of *BRCA2* expression and increased sensitivity to PARP inhibitors (Sup. Figure 3b).

In contrast to wild-type OVCAR8 cells, the *BRCA2* mutant clones displayed a significant increase in cell death upon treatment with ATR inhibitors alone or knockdown of MCM7 alone (Sup. Figure 3c). Consequently, the combination of both treatments resulted in only a minor additional increase in cell death. Similarly, we did not observe an additive or synergistic effect in terms of cGAS expression, micronuclei formation, or abnormal cell formation (Sup. Figure 3d, e). When the cells were pre-treated with XL413, RL5a, or Simvastatin, we observed a synergistic increase in cell death only in wild-type OVCAR8 cells, but not in the *BRCA2* mutant clones (Sup. Figure 3f). These findings suggest that the synergistic effect may be more efficient in cells with an intact HR pathway, as cells with a defective HR pathway are more sensitive to single treatments.

### Reducing the origin of replication pool in *CCNE1* amplified cells

Considering the association of cGAS expression with cell cycle processes and the role of Cyclin E1 in cancer development, we investigated whether high levels of *CCNE1* could amplify the synergistic effects observed in ovarian cancer cell lines when combining ATR inhibitors (ATRi) and replication origin reduction. To examine this, we utilized the immortalized fallopian tube cell line FT282, which expresses a doxycycline-inducible *CCNE1* vector.^31^

Importantly, after *MCM7* knockdown, cells expressing high levels of Cyclin E1 showed increased sensitivity to ATR inhibitors compared to cells expressing normal levels of Cyclin E1 (Figure 5a, b).

**Figure 5:**
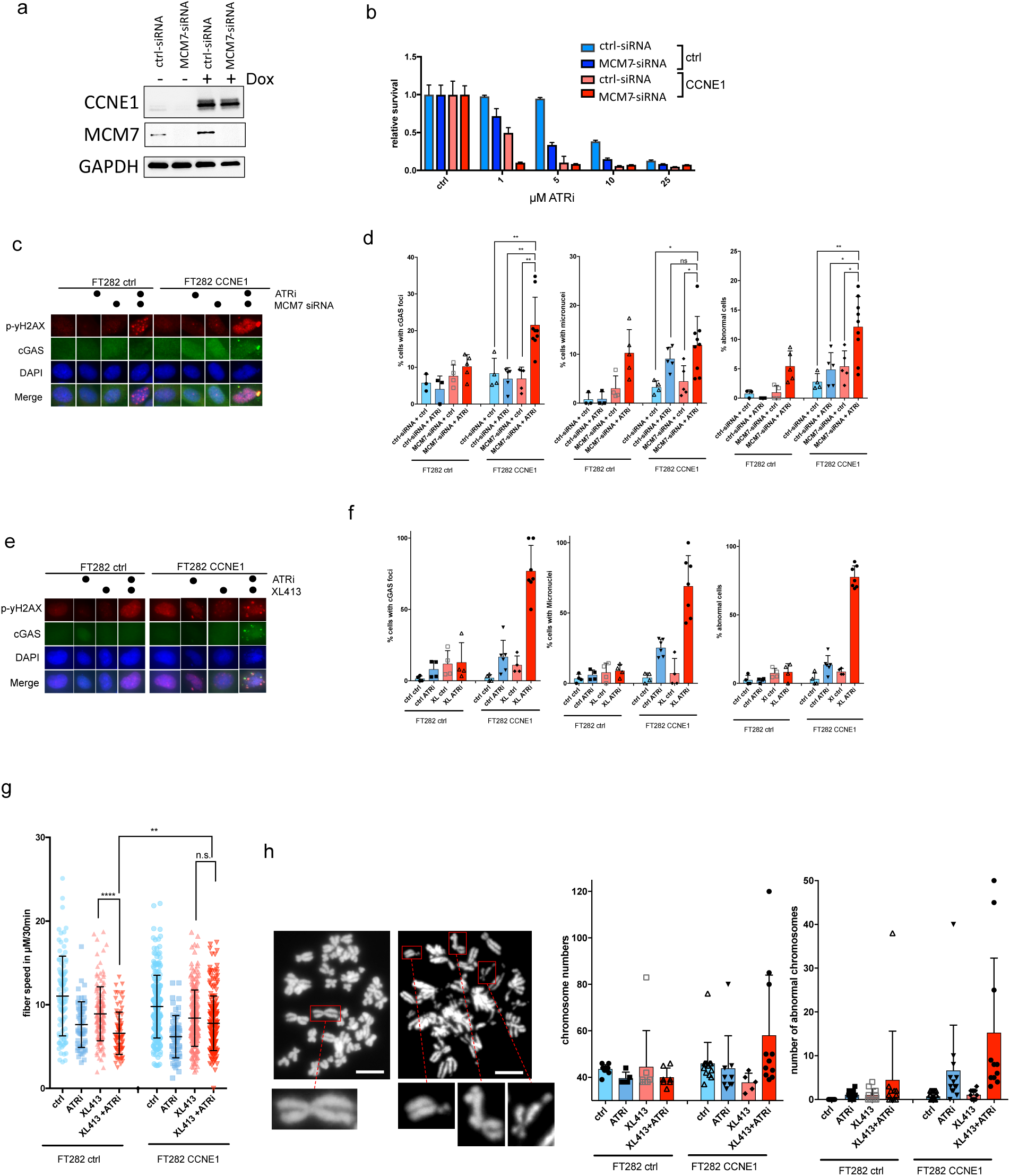
Reducing the origin of the replication pool in CCNE1 amplified cells. a) Western blot of FT282 cells with high and low Cyclin E1 expression treated with MCM7 or control siRNA. b) Relative survival analysis of FT282 cells with high and low Cyclin E1 expression treated with MCM7 or control siRNA followed by ATRi treatment. c and d) FT282 cells with high and low Cyclin E1 expression treated with MCM7 or control siRNA followed by ATRi treatment were analyzed by immunofluorescence (c, corresponding image) for percent of cells per image with cGAS foci, micronuclei or abnormal cells (d). e and f) FT282 cells expressing high and low Cyclin E1 expression treated with XL413 followed by ATRi treatment were analyzed by immunofluorescence (e, corresponding image)for percent of cells per image with cGAS foci, micronuclei or abnormal cells (f). g) DNA fiber spread assay as described in material and methods under different conditions described in e. h) Mitotic chromatin spread assay of conditions described in e. Left: representative image of spread chromosomes normal (left) and abnormal (right). Right: quantified are the chromosome numbers and abnormal chromosomes per spread nucleus. Scale bar 10 μm.

Activation of cGAS-STING pathway by cytosolic DNA has been shown to induce the Type I interferon response and downstream NF-KB signaling. Recently, it was demonstrated that immune activation by cGAS-STING is dependent on cell progression in a model of ionizing radiation. Given that Cyclin E1 overexpression also induces DNA double-strand breaks and forces cells through the cell cycle, we hypothesized that the combination of *MCM7* siRNA and ATRi might also activate the innate immune response.^28,29^

In Cyclin E1 overexpressing cells treated with ATRi and *MCM7* siRNA, we observed a significant increase in cGAS foci formation and the number of cells with abnormal nuclei, which was not observed in control cells (Figure 5c, d).

We then investigated whether the combination of XL413 and ATR inhibitors (ATRi) could also increase genomic instability. In FT282 cells overexpressing Cyclin E1, we observed a strong combinational effect on cGAS expression, micronuclei formation, and abnormal cell formation, whereas no such effect was observed in control FT282 cells (Figure 5e, f)

Furthermore, we analyzed replication speed in FT282 control cells and Cyclin E1 overexpressing cells. As expected, we observed a decreased replication speed in both control and Cyclin E1 overexpressing cells when treated with ATRi, indicating increased replication origin firing (Figure 5g). Interestingly, this effect was abrogated in Cyclin E1 overexpressing cells when co-treated with XL413, but not in control cells. This suggests that the effects of Cyclin E1 overexpression on replication origins, combined with the inhibition of replication origin firing, can reduce the effects of ATRi on replication activation. This is significant because replication speed and the distance between activated replication origins are directly linked. Cells with a faster replication speed have a longer distance between replication origins, while cells with more activated replication origins have a shorter distance between them, allowing them to tolerate a slower replication speed to replicate within the same timeframe.

Given the effects on replication, we expected changes in the size and shape of the nuclei in treated cells. While we observed a reduction in nuclear area and compactness in control cells treated with ATRi, we did not observe this effect in Cyclin E1 overexpressing cells (Sup. Figure 4a, b). When analyzing chromosome changes using mitotic spread assay, we observed a strong effect on chromosome number as well as on number of abnormal chromosomes in the Cyclin E1 overexpressing cells treated with XL413 and ATRi (Figure 5h).

In summary, we here show that the aberrant expression of Cyclin E1 amplifies the effects on genomic instability and innate immune activation of combined reduced replication activation and ATRi.

### Immune Effects on Cyclin E1 overexpressing cells

Finally, we investigated if the combination treatment of XL413 and ATRi could enhance the innate immune response in Cyclin E1 overexpressing cells. Control cells treated with XL413/ATRi showed no effect; however, cells overexpressing Cyclin E1 exhibited strong phosphorylation of STAT1 (Y701) and p65 (NF-KB) (Figure 6a). The increased phosphorylation was accompanied by higher mRNA expression of downstream target genes, including *CCL5*, *ISG15, CXCL10, IL6,* and *IFI44* (Figure 6b). Additionally, we observed an increased release of CCL5 into the media (Figure 6c).

**Figure 6:**
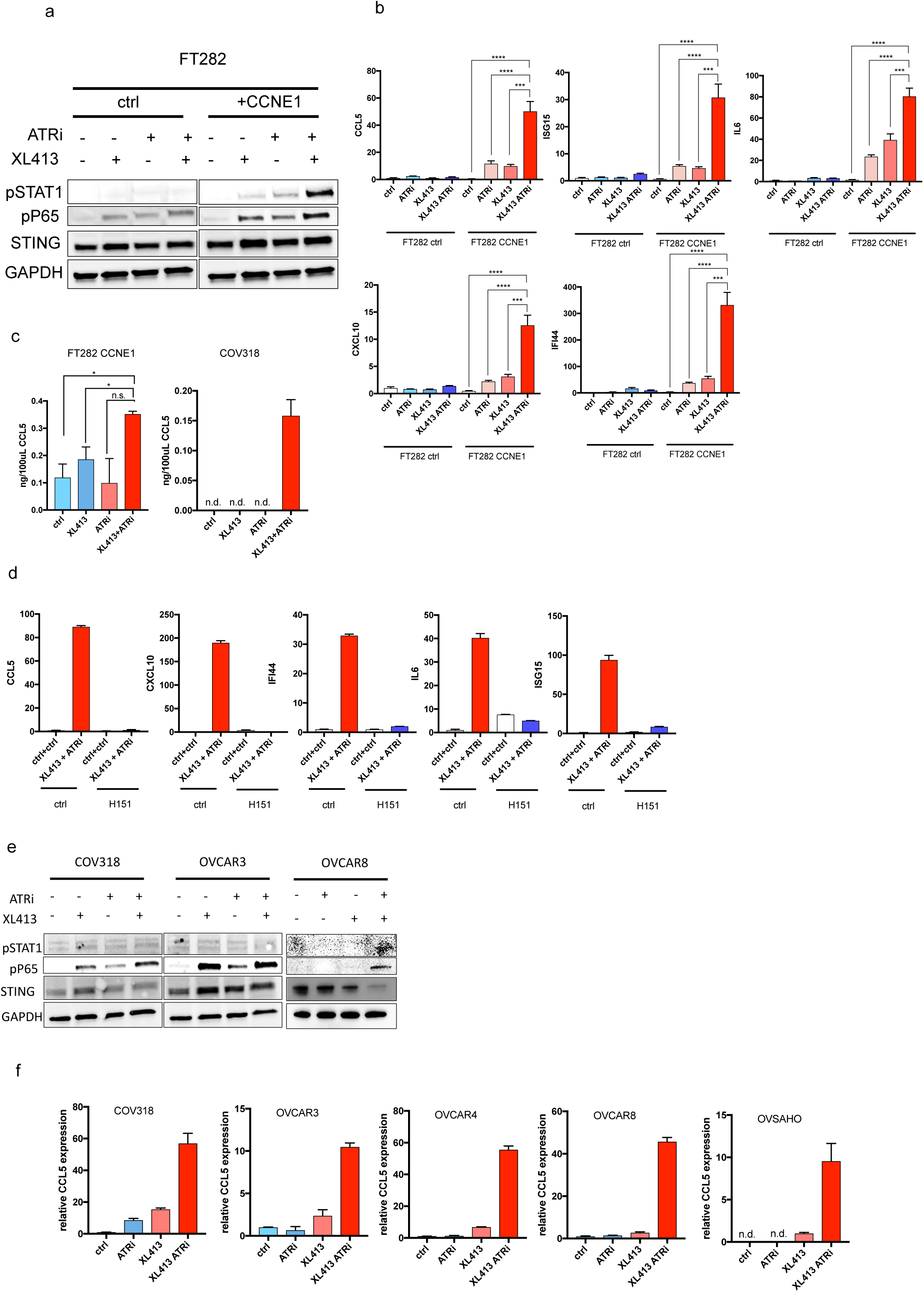
Immune effects on CCNE1 overexpressing cells. a) Western blot analysis of FT282 control and CCNE1 overexpressing cells treated with XL413 and ATRi alone or in combination. b) RNA expression of cells in (a) was analyzed by real time PCR for CCL5, ISG15, IL6, CXCL10 and IFI44. Depicted are fold changes in relation to the control sample. c) CCL5 expression measured by ELISA from the supernatant of different cell lines treated with XL413 and ATRi alone or in combination. d) Real time PCR of FT282 cells overexpressing CCNE1 treated with XL413 followed with the treatment of ATRi alone or in combination the STING inhibitor H151. e) Western blot analysis of COV318, OVCAR3, and OVCAR8 cells after the treatment with ATRi and XL413 alone or in combination. f) Real time PCR of CCL5 expression in multiple cell lines after XL413 treatment alone or in combination with ATRi.

To further validate the involvement of the STING pathway, we used the STING1 inhibitor H151, which covalently binds to STING1, to inhibit the STING signaling. Our data demonstrate that the inhibition of STING1 abrogates the downstream innate immune response in Cyclin E1 overexpressing cells (Figure 6d).

Furthermore, we tested multiple ovarian cancer cell lines and found that the combinatory treatment led to an increase in phosphorylated P65 in multiple cell lines (Figure 6e). The downstream effect on *CCL5* gene expression was also observed in multiple cell lines (Figure 6e). These findings suggest that combining XL413 and ATRi may be beneficial in inducing an innate immune response, not only in *CCNE1* amplified tumors, but in a wide range of tumor cells, potentially aiding in the recruitment of cytotoxic T-cells to the tumor.

## Discussion

In our study, we discovered that the reduction of replication origins or the inhibition of their activation can sensitize cancer cells to ATRi by inducing intolerable levels of genomic instability. The strong differences in the synergistic effects observed between cell lines could be attributed to varying levels of endogenous replication origin loading. The most pronounced synergistic effects were observed in cells with high Cyclin E1 levels due to overexpression or amplification. Cyclin E1 is a regulator of the cell cycle that controls the transition from the G1 phase to the S phase. Aberrant expression of Cyclin E1 leads to a shortened G1 phase, during which cells are unable to acquire the necessary resources for genome duplication in the subsequent S phase. This has two major consequences: First, cells fail to accumulate an adequate amount of nucleotides required for DNA replication, and second, fewer pre-replication complexes are loaded onto the genome.

The pre-replication complex generation is restricted to the G1 phase to reduce the chance of re-replication and amplification during the S and G2 phase. It is generated by sequential loading of the origin recognition complex (ORC), licensing factors CDT1and CDC6, and the MCM helicase complex onto chromatin.^21,32^

A reduced number of loaded replication complexes usually does not present a problem, since cells load replication origins in abundance, using only 10% of loaded origins and keeping the rest as reserve in case replication stress. In addition to the indirect effects on origin loading through G1 phase shortening, Cyclin E1 has been shown to directly facilitate MCM2-7 loading independently of its interaction partner CDK2. Interestingly, an aberrant Cyclin E1 expression impairs DNA replication by interfering with the pre-replication complex assembly during G1-phase resulting in a reduced number of licensed origins during S-phase.^13,30^ This reduced number of licensed origins subsequently leads to an increased distance between forks, thus increasing the average replicon size, resulting in unfinished chromosome replication, before the cell enters mitosis.^33,34^ As a consequence, the replicated chromosomes undergo breakage, resulting in a higher frequency of double-stranded DNA breaks and karyotypic abnormalities. Furthermore, it was shown that the increased replication initiation, due to Cyclin E1 overexpression, leads to replication stress that results from conflicts between replication and transcription, called transcription-induced replication stress.^35^

In our study, we found that the changes in replication origin loading can be exploited by decreasing the levels of MCM proteins through siRNA or small molecule inhibitors. Although reducing the MCM pool can result in genomic instability, it is challenging to affect normal healthy cells because they use only a small fraction (10%) of their replication origin reservoir for genome duplication.

Here we combined the reduction of replication origins with ATR inhibition because ATR interacts at different levels with the effects of changes in replication. Replication stress caused by incomplete replication or collapsing replication forks activates the ATR pathway leading to a cell cycle arrest. Without ATR, the cell continues to replicate and thereby accumulating genomic fractures that lead to genomic instability. ATR kinase inhibition has been also shown to induce unscheduled origin firing and reduces replication fork velocity.^21,36,37^ It was further shown that the ATRi induced origin firing is mediated by CDC7 kinase through phosphorylation on GINS.^21^ It is not clear how this translates to endogenous replication stress and DNA damage in the case of overexpressing Cyclin E1 cells.

Recently the group of Corrado Santocanale identified that the loss of the ATR activator ETAA1 reduces the sensitivity to CDC7i using a CRISPR-CAS9 genome wide screen.^24^ They further showed that the CDC7i XL413 induces ATR activation mainly through ETAA1, and that its inhibition leads consequently to origin activation that drives into premature mitosis. Our data agree with the data from Rainey et al, but further show that the underlying genetics is an important consideration for this combination. Specifically, *CCNE1* amplified or overexpressing tumors might be most responsive to this combination, whereas no synergistic effects were observed in *BRCA2* mutated cancer cells. We also show that it is possible to directly reduce the MCM pool using compounds like RL5 or Simvastatin in combination with ATRi.

Effective combinations must be further tested in pre-clinical and clinical models.

Statins like Simvastatin are widely used as lipid lowering medications in clinical routine and repurposing such drugs for other purposes like to further sensitize high-grade ovarian cancer to ATR inhibition as described here might be particularly appealing, due to their well described pharmacodynamic and safety profile.

Like Rainey et al. we also observed that the combination with inhibitors against ATR and CDC7 leads to grape or abnormal shaped nuclei which are associated with mitotic defects and containing variable amounts of DNA.^24^ Possibly, for the optimal efficacy, the loss of the S-G2-M checkpoint combined with the reduced replication origins are required together with additional cell cycle factors that push the cell through the cell cycle such as Cyclin E1. Beside the grape shaped structures we also observed a strong increase in micronuclei. The laboratory of Roger Greenberg has found that micronuclei are induced by DNA-double strand breaks that are pushed through mitosis.^28^ They further found that this activation is responsible for the activation of the pattern-recognition receptor cyclic GMP-AMP synthase (cGAS). They showed that the anti-tumor inflammatory signaling can be prevented through prolonged G2 arrest.^29^ In line with these findings we found a strong increase in micronuclei formation that is associated with an increase in cGAS-STING signaling. The mis-segregation of chromosomes during mitosis has been shown to lead to the formation of abnormal nuclei and micronuclei. Approximately 50% of micronuclei are unstable, releasing double stranded DNA to the cytoplasm allowing it to be recognized by cGAS.^38,39^ In contrast to the previous finding that showed that STING loss did not eliminate the irradiation induced inflammation, in our approach we were able to completely abrogate the inflammatory stimulated gene expression.^28^ It was further shown *in* vivo that the inflammatory response following IR was enhanced through the combined loss of G1 and G2 cell cycle checkpoint through *TP53* mutation and ATRi, respectively.^40^ In HGSOC *TP53* is ubiquitously lost due to mutations and represents one of the earliest events during tumor development.^41^ The fallopian tube cell line FT282 was immortalized using hTERT and an shRNA targeting *P53*.^31^ In this model ATRi was not sufficient to induce micronuclei and an innate immune response. An additional reduction of replication origins or an overexpression of Cyclin E1 was needed to increase micronuclei formation. To further induce a strong innate immune response, the reduction of replication origins and an overexpression of Cyclin E1 was necessary, indicating that this combination might specifically target tumor cells. Interestingly, in cancer cell lines we observe this effect independently of Cyclin E1 expression, indicating that other factors might also contribute to this effect. A current clinical challenge is that in contrast to melanoma, non-small cell lung cancer and other cancer types, single-agent antibodies targeting CTLA-4, PD-L1 or PD-1 only resulted in very modest responses in HGSOC patients.^42–44^ Despite those challenges it is known that HGSOC patients with T-cell-rich tumors experience longer overall survival.^45^ Nevertheless, since HGSOC is a copy number driven tumor that has only few recurrent mutations, tumors with high T-cell levels are rare.^45^ While *BRCA1/2* mutated tumors show a higher neoantigen load, and PD-1/PD-L1 expression that is not true for HR-proficient tumors such as tumors with a *CCNE1* amplification.^46,47^ Therefore, it is a current clinical challenge to increase T-cell infiltration in HGSOCs.

Our findings demonstrate that the combination of a CDC7 and an ATR inhibitor represents a promising approach to increase the effectiveness of treating Cyclin E1 overexpressing tumors while also activating the innate immune system. Further pre-clinical and clinical studies will be needed to assess the potential of this combination therapy in future patient treatment.

## Methods

### Bioinformatics

The data from the TCGA Ovarian cancer cohort were extracted and downloaded from the XENA portal of the University of California, Santa Cruz (http://xena.ucsc.edu/) and the cBio-Portal website ^48^. Cell line expression data and drug response data were downloaded from the https://depmap.org/website.^49^ CTL data were extracted from the TIDE website (http://tide.dfci.harvard.edu/).^50^ Synergies were calculated using the SynergyFinder 2.0 R package and website.^51^ using Data were analyzed with the R programming language and the program GraphPad Prism. R packages included complex heatmap.^52^

### Western blot and antibodies

For western blotting cells were lysed in RIPA buffer supplemented with protease inhibitors, for 30 min on ice. Proteins content was quantified by Bradford assay (Bio-Rad). 20 µg proteins were analyzed through a 4-15% gradient SDS-PAGE before being transferred to the PVDF membrane with the TurboBlot system (Bio-Rad). The membrane was blocked with 5% (w/v) BSA in TBS-tween for one hour at room temperature before incubating in primary antibody overnight at 4°C. All antibodies are listed in Supplementary Table S1. After washing three times with TBS-tween the membrane was incubated either with an anti-mouse or an anti-rabbit HRP-conjugated secondary antibody for 1 hour. Protein bands were detected using Clarity Chemiluminescent HRP Antibody Detection Reagent (Bio-Rad) and visualized with a chemiluminescence reader (Fusion SL) imaging system.

### Cell culture and cell lines

The cell lines FT282, KURAMOCHI and OVSAHO cells were a gift from Ronny Drapkin (UPenn, Philadelphia, USA). The establishment of the FT282 cell line has been previously described ^31^. The ovarian cancer cell lines (OVCAR8, OVCAR3, HEYA8 and SKOV3) were obtained from American Type Culture Collection (ATCC, Manassas, VA). All cancer cell lines were cultured in DMEM F12 (Invitrogen, Carlsbad, CA) supplemented with 10% fetal bovine serum (FBS, Atlanta Biologicals) and 1% penicillin/streptomycin (Invitrogen). All cells were grown inside an incubator maintaining 37°C and a 5% CO2-containing atmosphere.

### Short-interfering RNA transfection

A pool of three short-interfering RNA (siRNA) duplexes of the trilencer-27 siRNA (Origene) was used to downregulate the MCM7 protein expression. As a negative control an unspecific scrambled trilencer-27 siRNA was used. Twenty-four hours after seeding 1 × 10^5^ cells in six-well plates, the cells were transfected with the siRNA using Lipofectamin RNAiMAX (Invitrogen, Karlsruhe, Germany) together with 10 nM siRNA duplex per well as described in the instructions.

### Cell survival

Cell growth and survival was measured using a CellTiter 96® Non-Radioactive Cell Proliferation Assay at an absorbance of 570 nm using (Promega, Mannheim, Germany). Each assay was performed in triplicates and repeated at least 3 times. Data are presented by means ± SD. Statistical and significant differences were determined by ANOVA with post-hoc analysis. Cells were additionally stained with crystal violet to count the remaining attached cells.

### Colony form

500 cells were seeded on 12-well plates and cultured for two weeks. Cells were then washed with PBS and stained with the Crystal Violet buffer (Crystal Violet 0.05% w/v, Formaldehyde 1%, 10X PBS (1X), Methanol 1%) for 20 min. Cells were washed 3 times with water and air dried.

### Immunofluorescence

Cells were grown overnight on coverslips in 96-well Cell Imaging Plates (Eppendorf). After washing, cells were fixed in 4% (v/v) paraformaldehyde in PBS for 20 minutes at room temperature. Cells were permeabilized and blocked with super-block buffer (Thermo Scientific) supplemented with 0.3% Triton X-100 for 10 minutes. Cells were incubated with primary antibodies for 2 hours at room temperature or overnight in the cold room. Following three washing steps the secondary antibody conjugated to Alexa Fluor Dyes (Molecular Probes) was incubated for 45 minutes at room temperature. Nuclei were stained with DAPI for 3 minutes. After three washing steps the cells were mounted with Flouromount-G (Sigma-Aldrich) prior to microscopy using a Zeiss Axio Observer inverted microscope with ×40 magnification.

### Fiber spread

1 × 10^5^ cells per well were plated in each six well. 72 hours after treatment, cells were labeled with CIdU (38µM) for 30 minutes, washed 3 times before medium containing 250 µM IdU was added for 30 minutes. Cells were trypsenized, washed and re-suspended in 30 µl PBS. Directly on a glass slide 2.5 µl cell suspension was mixed with 7.5 µl of lysis buffer (200 mM TrisHCl pH7.4, 50 mM EDTA, 0.5% SDS). Slides were tilted manually 45⁰ to allow the drop to run slowly down the slide and air dried. The DNA was fixed in cold methanol/acetic acid (3:1 o/n) for 5 minutes. Rehydrated slides were denatured in 2.5M HCl for one hour. After washing the slides were blocked for 30 min in Superblock solution following the incubation with the primary antibodies mouse anti-BrdU/IdU (1:100, BD Bioscscience) and rat anti-BRDU/CIdU (1:200, Abcam) for overnight at 4C. After washing, slides were incubated with secondary fluorescent antibodies (anti-mouse alexa 488, 1:300, Molecular Probes, anti-rat Cy3, 1:300 Jackson Immuno Research) diluted in blocking solution for one hour. Slides were washed and air dried before mounted with coverslip and 20 µl Antifade Gold (Invitorgen). Fibers were examined using a DMRE, upright, Light Microscope Leica fluorescence microscope with a 100×oil immersion objective and analyzed using ImageJ software.

### Cell cycle analysis

Cells from one six-well were trypsinized, washed with cold PBS and resuspended in 300µl cold PBS. 700µl 100% ice cold ethanol was added dropwise and samples were incubated at −20C for at least 24 hours. Cells were washed in PBS and then incubated for 30 minutes in 300µl DAPI/Triton X-100 PBS Solution (0.1% (v/v) Triton X-100, 1µg/ml DAPI). Cells were analyzed in parameters 355-450nm in linear mode.

### Chromosome spread

1×10^5^ cells were grown for different experimental conditions in a 6 well plate. After different treatment conditions, a final concentration of 0.02µg/ml Colcemid was added to the cells for 90 minutes at 37C. Cells were harvested by trypsinization and washed 3 times. Cells were resuspended in 1ml of hypotonic solution (0.075 M KCl) and incubated for 15 minutes at 37C. Cells were centrifuged and fixed in 300µl fixing solution (3:1 Methanol:Acetic Acid) and incubated for 30 minutes. This step was repeated 2 times for 10 minutes. 20µl cells were dropped from about 10cm high onto glass microscope slides and run through a flame of a bunsen burner. Dried slides were stained with DAPI (1µg/ml) for 5 minutes, air dried and fluoromount with a fighting cover slide. Chromosomes were examined using a 100x oil immersion lens with a Light Microscope Leica fluorescence microscope.

### Real time PCR

RNA was isolated using the RNeasy Plus Mini Kit (Qiagen) as described in the manufacturers protocol and measured using a nanodrop apparatus. 0.5µg were transcribed to cDNA using the RT2 Easy First Strand Kit (Qiagen). Real time PCR was performed using the POWRUP SYBR MASTER MIX (Life Technologies) and listed primers. We used the QuantStudio Real-Time PCR systems (Thermofisher) to quantify ct-values. The relative gene expression was calculated using the delta-delta -ct value.

### Crispr CAS9

OVCAR8 cells were seeded in a 24-well plate 24h prior to transfection to 80% confluency. 3pmol of the Cas9 Nuclease and 3.9pmol of each *BRCA2* sgRNAs (Sequences: UCUACCUGACCAAUCGA; UAGCACGCAUUCACAUA, were diluted in 25µL Opti-MEM media together with 1µL Lipofectamine Cas9 Plus reagent (Synthego CRISPR system). 1.5µL of the Lipofectamine CRISPRMAX transfection reagent diluted in 25µL Opti-MEM media were then mixed with the diluted nucleic acid dilution and mixed gently. The nucleic acid/transfection reagent solution was incubated at room temperature for 5 minutes before added to the cells. The cells were incubated for 72 hours before seeding single cells to 96-well plates. Grown clones were expanded and analyzed by Western blot for the loss of *BRCA2* expression. Potential clones were sequenced to confirm cleaved sequences. We used 2 different clones for this study.

### Statistics

All correlation values were calculated with the Pearson correlation coefficient. P-values were calculated with an unpaired two-sided t-test. *=p<0.05, **=p<0.01, ***=p<0.001, ****=p<0.0001

## Acknowledgements

We thank members of the Marmé/Doberstein lab and the department of gynecology Mannheim for fruitful discussions and comments. We thank Stefanie Gaiser of the Marmé/Doberstein lab for her technical support. This work was supported by the AstraZeneka iEvsion Grant.

## Author contributions

K.D.: Conceptualization, methodology, validation, formal analysis, investigation, data curation, writing—original draft, software, visualization, supervision, project administration, and funding acquisition. J. P.: Methodology and data curation. S.B. and B.T.: Investigation and resources. M.S.: Investigation, resources, and funding acquisition. F.M.: Investigation, resources, supervision, project administration, and funding acquisition.

