## Supplementary Information for "Inhibition of Replication Origins and ATR Synergistically Activates the Innate Immune System in Cancer Cells"

### Supplementary figures

Supplementary Figure 1

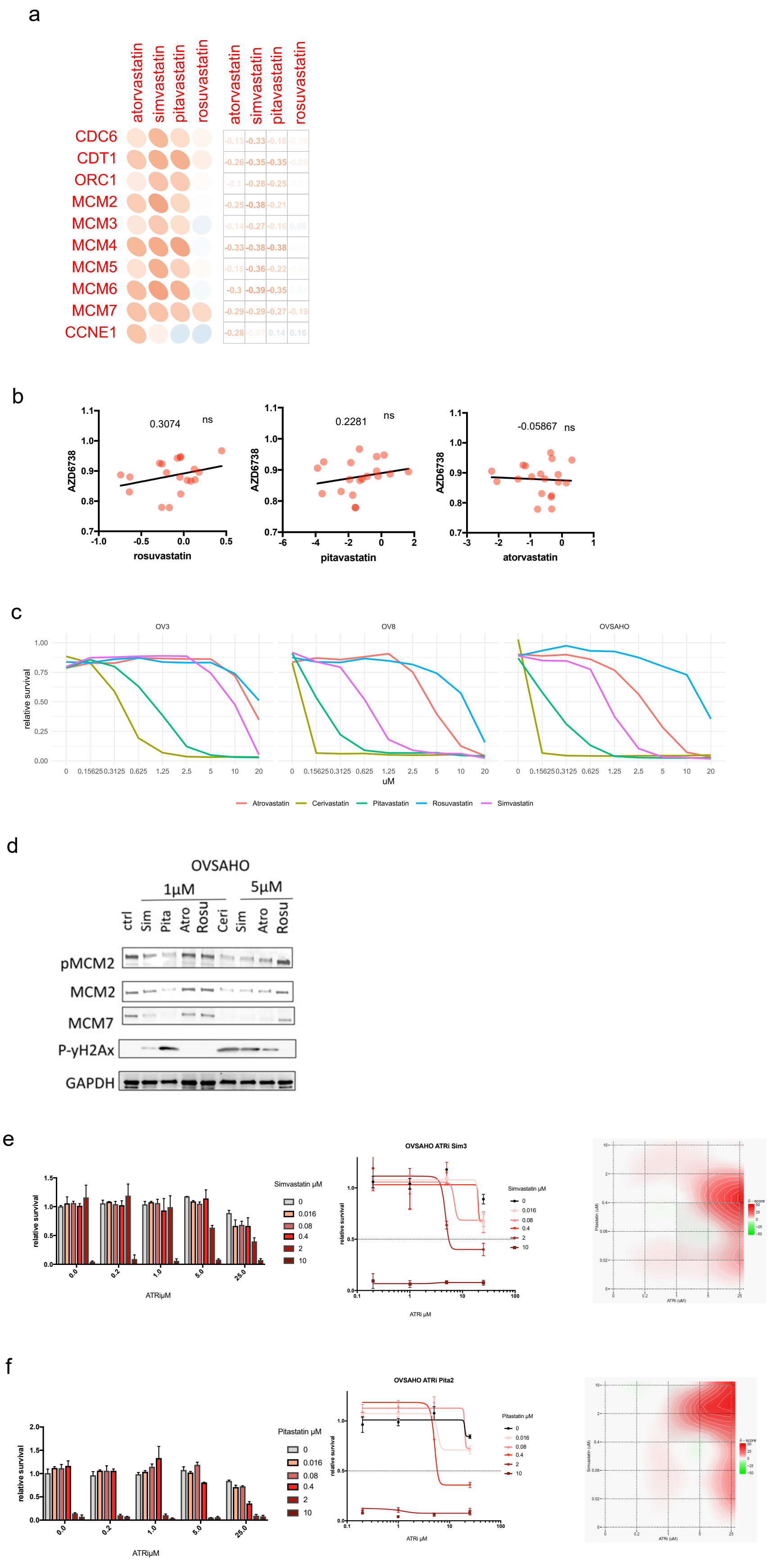

**Sup. Figure 1: Statins can reduce the origin Pool.** a) Correlation matrix of the expression of origin of replication genes with the AUC of Atorvastatin, Simvastatin, Pitavastatin and Rosuvastain in ovarian cancer cell lines from the cancer cell line encyclopedia and the depmap datasets. b) Diagram of AUC of AZD6738 and different statins in ovarian cancer cell lines from the depmap datasets. c) Survival curve of OVCAR3, OVCAR8 and OVSAHO cells treated with different concentrations of statins. d) Western blot protein expression in OVSAHO cells after the treatment with 1µM or 5µM of different statins. e and f) Survival analysis of OVSAHO cells treated with different concentrations of simvastatin (e) or pitavastatin (f) followed by ATRi treatment. On the right is the synergy profile depicted.

Supplementary Figure 2

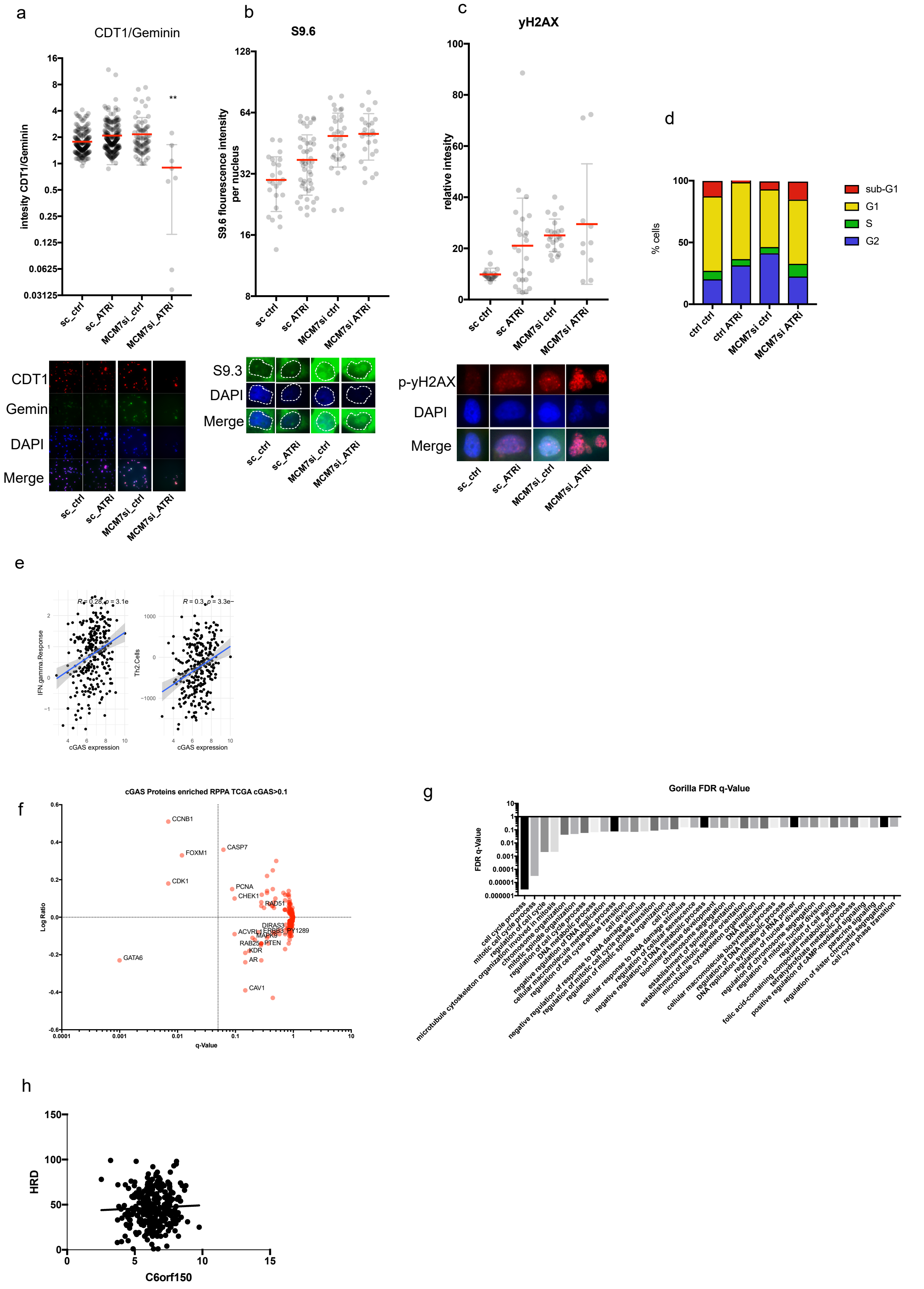

**Sup. Figure 2: Reduction of the replication origins leads to chromosomal abnormalities.** a) Immunofluorescence analysis of CDT1 (red) and Geminin (green) in OVCAR8 cells treated with control or MCM7 siRNA followed by ATRi treatment. b) Immunofluorescence analysis of nuclear DNA/RNA hybrids by measuring S9.3 antibody fluorescence intensity (green) in the nucleus of OVCAR8 cells. c) Phospho-yH2AX nucleus intensity measured in OVCAR8 cells. d) cell cycle analysis in OVCAR8 cells. e) Diagrams of INF-gamma response (left) or TH2-cells (right) and cGAS RNA expression (C6orf150). f) Vulcano blot of enriched proteins measured by RPPA in ovarian cancer tumors expressing high levels of cGAS. g) Waterfall plot of the FDR-qValue of enriched pathways in ovarian cancers expressing high cGAS levels. h) Diagram showing HRD score and cGAS RNA expression in the TCGA ovarian cancer dataset.

Supplementary Figure 3

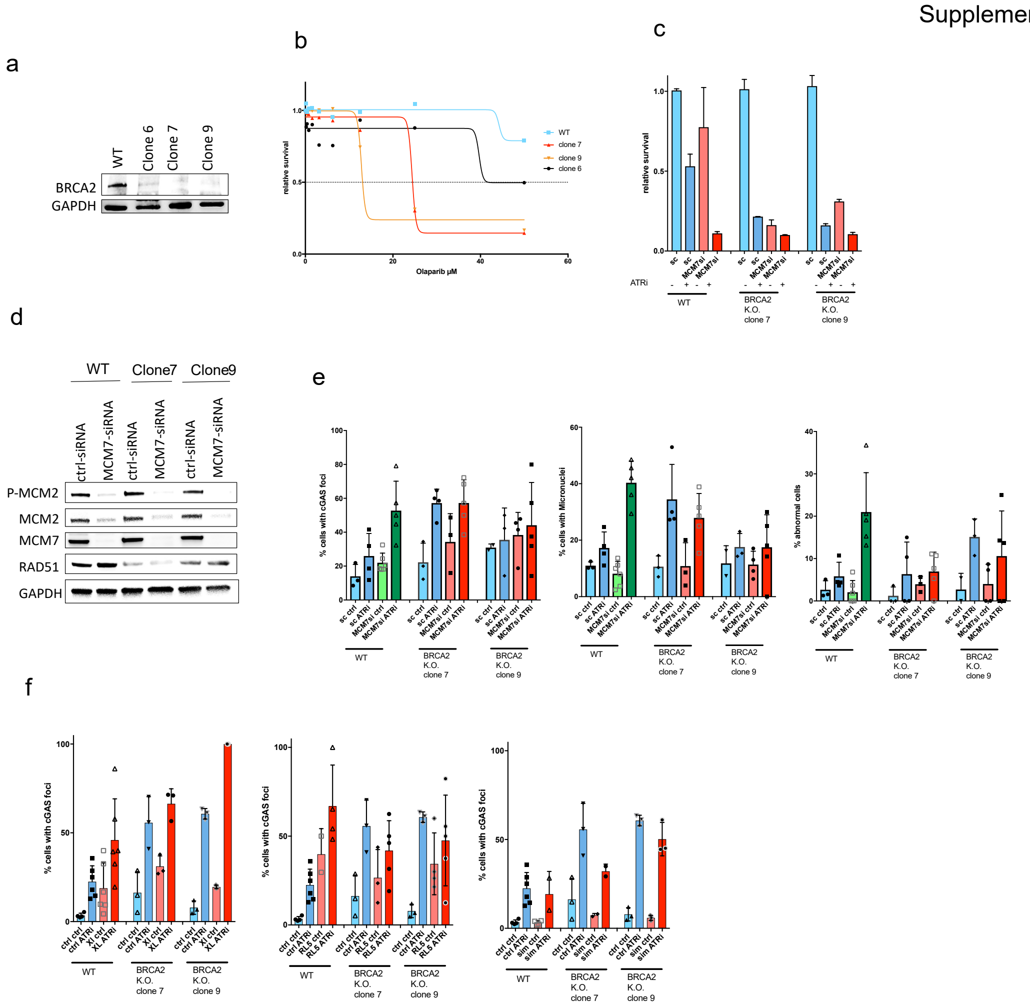

**Sup. Figure 3: Reducing the replication origin pool in BRCA2 mutated cells.** a) Western blot analysis of BRCA2 CRISPR CAS9 knockout clones. b) Sensitivity to Olaparib measured in different clones. c) survival analysis of WT OVCAR8, clone 7 and clone 9 following the treatment with MCM7 siRNA and ATRi. d) Western blot analysis from experiment c. e) Percent of cells with cGAS foci, micronuclei and abnormal cells analyzed in cells from c. f) Percent cGAS positive cells were analyzed in WT OVCAR8, clone 7 and clone 9 after the treatment with XL413 (left), RL5 (middle) or Simvastatin (right) followed by the treatment with ATRi.

Supplementary Figure 4

a

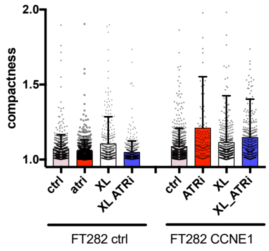


b

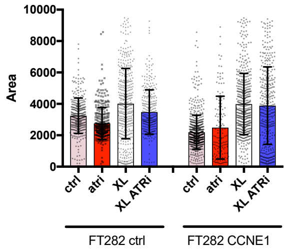

**Sup Figure 4: Reducing the origin of the replication pool in CCNE1 amplified cells.** a and b) compactness (a) and Area (b) of nuclei measured in FT282 cells after treatment with XL413 alone or in combination with ATRi.
